# Epigenetic control of tetrapyrrole biosynthesis by ^m4^C DNA methylation in a cyanobacterium

**DOI:** 10.1101/2024.08.20.608618

**Authors:** Nils Schmidt, Nils Stappert, Kaori Nimura-Matsune, Satoru Watanabe, Roman Sobotka, Martin Hagemann, Wolfgang R. Hess

## Abstract

Epigenetic DNA modifications are pivotal in eukaryotic gene expression, but their regulatory significance in bacteria is less understood. In *Synechocystis* 6803, the DNA methyltransferase M.Ssp6803II modifies the first cytosine in the GGCC motif, forming N4-methylcytosine (GG^m4^CC). Deleting the *sll0729* gene (Δ*sll0729*) caused a bluish phenotype due to reduced chlorophyll levels, which was reversed by suppressor mutations. Re-sequencing of seven suppressor clones revealed a common GGCC to GGTC mutation in the *slr1790* promoter’s discriminator sequence, encoding protoporphyrinogen IX oxidase, HemJ, crucial for tetrapyrrole biosynthesis. Transcriptomic and qPCR analyses indicated aberrant *slr1790* expression in Δ*sll0729* mutants. This aberration led to the accumulation of coproporphyrin III and protoporphyrin IX, indicative of impaired HemJ activity. To confirm the importance of DNA methylation in *hemJ* expression, native and mutated *hemJ* promoter variants were introduced into the wild type, followed by *sll0729* deletion. The *sll0729* deletion segregated in strains with the GGTC motif in the *hemJ* promoter, resulting in wild-type-like pigmentation, whereas freshly prepared Δ*sll0729* mutants with the native *hemJ* promoter exhibited the bluish phenotype. These findings demonstrate that *hemJ* is tightly regulated in *Synechocystis* and that N4-methylcytosine is essential for proper *hemJ* expression.

## 1. Introduction

DNA base methylation occurs in all kingdoms of life. It affects DNA replication, cell cycle control, DNA mismatch repair, DNA-protein interactions, phenotypic heterogeneity among populations, gene expression, and recognition of intruder DNA.^1^ The most common types of methylated bases comprise the endocyclic 5-methylcytosine (^m5^C), the exocyclic N-6-methyladenine (^m6^A), and N4-methylcytosine (^m4^C). Newly methylated bases are established by DNA methyltransferases (MTases) that recognize specific DNA motifs.^2^ These modifications do not impair with Watson-Crick base pairing but protrude to the major groove of DNA exposing the methyl group to DNA binding proteins or the transcriptional machinery.^2^ Bacterial MTases are commonly associated with restriction endonucleases forming restriction-modification (R-M) systems.^3,4^

However, MTases not associated with a cognate restriction enzyme, named ‘orphan’ MTases, are also present. Global analyses of the DNA methylome of 230 different bacteria and archaea revealed such orphan MTases in 48% of the examined organisms.^5^ The most prominent examples of ‘orphan’ MTases in prokaryotes are the ^m5^C MTase Dcm and the ^m6^A MTases Dam and Ccrm.^6–8^ In eukaryotes and prokaryotes, ^m5^C^9–11^ as well as ^m6^A modifications can alter gene expression if associated with promoter sequences.^12,13^ CpG dinucleotides carrying a ^m5^C are most frequently observed in eukaryotic promoter regions,^10^ whereas e.g. the *pap*-operon of *E. coli* is regulated by ON and OFF states caused by the change in methylation of adenines in the promoter region.^14^ The effects of ^m4^C methylation in prokaryotes, however, has remained largely unclear. Changes in transcription and a change in pathogenicity were shown for *Leptospira interrogans*.^15^ But, how the occurrence of ^m4^C in promoter sequences can affect gene expression has still to be elucidated.

Cyanobacteria are the only prokaryotes conducting oxygenic photosynthesis making them ecologically relevant. Their unique physiology and redox chemistry have generated interest for producing various chemicals based on photosynthesis, using CO_2_ and solar energy.^16–18^ In the model cyanobacterium *Synechocystis* sp. PCC 6803 (*Synechocystis* 6803) five MTases have been identified. We previously showed that: i) the CGATCG sequence motifs in *Synechocystis* 6803 are double-methylated (^5m^CG^6m^ATCG) by the enzymes M.Ssp6803I and M.Ssp6803III; ii) these motifs are abundant within the repeat-spacer arrays of a subtype I-D CRISPR-Cas system; iii) conjugation efficiency decreased by 50% in cells lacking ^5m^C methylation due to deletion of gene *slr0214* encoding M.Ssp6803I.^19^ These results pointed at an important role of DNA methylation in *Synechocystis* 6803.

M.Ssp6803II, the DNA methyltransferase encoded by gene *sll0729* is responsible for the addition of an exocyclic methyl group to the internal cytosine in the motif GGCC, which occurs 38,512 times in the *Synechocystis* 6803 genome.^20^ Mutant strains with a deletion of gene *sll0729* (Δ*sll0729*) developed a pronounced phenotype, including slower growth, smaller cell diameter, lowered DNA contents, and a bluish pigmentation caused by a lowered chlorophyll/phycocyanin ratio,^21^ whereas complementation by *sll0729* restored the phenotype of the wild type (WT). We demonstrated that altered transcript levels of two genes, *sll0470* and *sll1526*, in Δ*sll0729* were caused by the lacking ^m4^C methylation in GGCC elements located in the respective −35 promoter elements.^21^ The results suggested the finding that ^m4^C plays some essential role in *Synechocystis* 6803. After longer cultivation times, suppressor mutants with a pigmentation resembling WT appeared frequently. These suppressor mutants were still ^m4^C methylation negative, but the nature of these mutations remained enigmatic.^21^

In this work we have addressed the molecular basis of the suppressor mutants. By resequencing analysis we found that all lines shared a single point mutation in one GGCC site recognized by M.Ssp6803II for ^m4^C methylation, which was changed in all sequenced suppressors to GGTC. However, this point mutation is not located in a coding region, but upstream of gene *slr1790* encoding protoporphyrinogen IX oxidase (HemJ), a key enzyme of tetrapyrrole biosynthesis.^22,23^ Transcriptomic analyses revealed a lowered mRNA level for *slr1790* in Δ*sll0729*, while levels similar to WT were reached in the suppressor mutants. In a series of reconstitution experiments we demonstrate that the non-balanced *hemJ* expression due to the non-methylated promoter sequence led to a dramatic over-accumulation of coproporphyrin III (CoPP) and protoporphyrin IX (PPIX), which is typical for the inhibited HemJ enzymatic activity (see below). In contrast, the accumulation of these metabolites became WT-like in the presence of the GGTC promoter variant. Overlay of the resequencing data with high-resolution data from the genome-wide mapping of transcriptional start sites suggested that the identified C-->T mutation is located within the discriminator area of the *slr1790* promoter. Thus, we show that N4 methylation of the first cytosine in the motif GGCC leading to GG^m4^CC is essential for the quantitative correct transcription initiation of the RNA polymerase and how a single nucleotide exchange of this cytosine in the promoter of *slr1790* can alter gene expression and affect the physiology of a photosynthetic bacterium. Our results underline the relevance of epigenetic effects linked to ^m4^C methylation.

## 2. Material and Methods

### 2.1. Bacterial strains and cultivation

*Synechocystis* sp. PCC 6803 substrain PCC-M^24^ was used in all experiments and served as wild type (WT). Axenic strains were maintained on agar plates with BG11 mineral medium at 30°C under constant illumination (30 µmol photons m^−2^ s^−1^; to slow down formation of suppressor clones). Transformants were selected and segregated on media containing 50 µg ml^−1^ kanamycin (Km; Sigma), 50 µg ml^−1^ spectinomycin (Spec; Sigma) or both. For physiological characterization axenic cultures of the different strains were grown photoautotrophically in BG11 medium, either under slight shaking in Erlenmeyer flasks at 30 µmol photons m^−2^ s^−1^, or aerated with atmospheric air in a photobioreactor (Multi-Cultivator MC 1000-OD-WW warm light, Photon Systems Instruments) at 30°C under continuous illumination of 40 µmol photons m^−2^ s^−1^. The appearance of suppressor clones was detected by measuring the absorption spectrum in between 400 and 750 nm (CARY 50 Bio, Varian). Contamination by heterotrophic bacteria was evaluated by spreading 0.2 ml culture aliquots on LB plates. The *E. coli* strain DH5α was used for routine DNA manipulations. *E. coli* was cultured in LB medium at 37°C. *Synechocystis* 6803 growth was recorded by measurements of optical density at 720 nm (OD_720_) either manually (shaking flask) or automatically (Multicultivator).

### 2.2. Resequencing analysis of suppressor lines

Five-hundred ng of genomic DNA was fragmented to an average length of 500 bp using a Covaris S2 sonication system (Covaris, Inc., MA, USA). Sequencing libraries were prepared using the NEBNext Ultra DNA Library Prep Kit for Illumina (New England Biolabs). Paired-end sequencing was carried out for 150 cycles using the Nextseq500 system (Illumina Inc., CA, USA) according to the manufacturer’s specifications. The sequencing reads were trimmed using the CLC Genomics Workbench ver. 9.5.4 (Qiagen) with the following parameters: Phred quality score > 30; removing the 5′ terminal 10 nt; and removing truncated reads <20 nt. Trimmed reads were mapped to the reference genome sequence and plasmids of *Synechocystis* 6803 (accession numbers: CP003265-CP003272) using CLC Genomics Workbench ver. 9.5.4 (Qiagen) with the following parameters: length fraction: 0.8, and similarity fraction: 0.9. To identify SNVs and indels, we used the filter settings as follows: minimum read depth for SNV/indel calling, 20; minimum read depth for the SNV calling, 10. Sequence reads were deposited to the DRA/SRA database with the accession numbers DRR585585–DRR585594, BioProject PRJDB18568.

### 2.3. Genetic engineering

The isolation of total DNA from *Synechocystis* 6803 was performed as previously described.^20^ The GG^m4^CC methylation was verified by restriction analyses using chromosomal DNA from *Synechocystis* 6803, the restriction endonucleases were used in a 10-fold excess and were incubated for at least 16 h at 37°C to ensure complete digestion. Synthetic primers were deduced from the genome sequence of *Synechocystis* 6803^25^ for the specific amplification of the promoter sequence of the *slr1790* gene, coding for the protoporphyrinogen IX oxidase HemJ. Mutagenesis PCR was used to mutate the GG^m4^CC methylation site of the promoter sequence to GGTC (see **Supplementary Table S4** for primer sequences). For this purpose, a DNA fragment containing the promoter region of *slr1790* was amplified by PCR and cloned into pJET1.2 (Thermo Scientific). The *aadA* gene, conferring Spec resistance from pUC4S (Pharmacia), was introduced into selected restriction sites. Verified constructs were transferred into *Synechocystis* 6803.^26^ For generation of M.Ssp6803II-deficient strains, the *sll0729* gene was deleted as described.^20^ A list of all *Synechocystis* 6803 strains used in the study is given in **Supplementary Table S2**.

To check the *slr1790* promoter activity, different promoter variants with single nucleotide substitutions of the internal cytosine in GGCC were tested. The nucleotide sequence of the used *slr1790* promoter ranging from −143 to +38 (TSS at +1) was fused to *luxAB* reporter genes and integrated into the genome. Promoter variants carrying the WT motif GGCC, GGTC, GGAC and GGGC were generated. For the integration into the genome the pILA vector^27^ was used.

### 2.4. RNA preparation and qRT-PCR

Cultures were grown under standard light conditions (50 µmol photons m^−2^ s^−1^) and 30°C and harvested via centrifugation (5,000 g, 20°C, 10 min) during exponential phase (OD 0.8). The pellet was resuspended in 1 ml PGTX,^28^ immediately frozen in liquid nitrogen and processed further as previously described.^29^ Relative amounts of RNA were quantified by qRT-PCR. Therefore, cDNA was prepared according to the manufacturer’s instructions using 600 ng of each RNA sample with the QuantiTect Reverse Transcription Kit (QIAGEN). qRT-PCR was performed using the FastGene^®^ qFYR Real-Time PCR System (NIPPON Genetics). PCR cycles for each run are described in **Supplementary Dataset S1** and **S2**. qPCRBIO SyGreen Blue Mix Separate-ROX was used and PCR reactions were performed according to the protocol (PCR biosystems). Primers (**Supplementary Table S4**) were synthesized by IDT. All reactions were measured as technical triplicates and *rnpA* was used as an endogenous control. The data was analyzed using the qFYR Analyzer Studio Software (Nippon Genetics) and relative quantification of RNA was calculated using the comparative Ct Method (ΔΔCt). The WT sample was used as the calibrator. The average of each technical triplicate measurement was displayed as a bar chart.

### 2.5. Pigment quantifications

The strains were precultured at 30 µmol photons m^−2^ s^−1^ constant light and 30°C. After seven days the BG11 medium was refreshed and the cultures were grown for another two days (OD_720_ 0.15 – 0.3). All following steps were performed at dark conditions. 2 ml of the culture was collected by centrifugation (4,000 g, 20°C, 5 min). Supernatant was completely removed and the pelleted cells were resuspended in 200 µl A. dest. (HPLC grade, Merck) followed by centrifugation (4,000 g, 20°C, 2 min). The supernatant was discarded and the pellet was resuspended in 100 µl 75% methanol (HPLC grade, Merck) and incubated for 15 min at 20°C. The sample was then centrifuged (4,000 g, 20°C, 2 min) and the supernatant collected. Remaining pellet was resuspended in 80% methanol, incubated for 15 min at 20°C and centrifuged again (4,000 g, 20°C, 2 min.) Supernatant was collected, combined with the supernatant from the first extraction and the resulting solution centrifuged (4,000 g, 20°C, 5 min). The extracted pigments were finally transferred into micro inserts for high performance liquid chromatography (HPLC) analysis, performed as described.^30^

### 2.6. Microscopy and determination of cell size

Strains were cultivated at 30 µmol photons m^−2^ s^−1^ constant light and 30°C for 10 days. Ten µl of these cultures were transferred on microscope slide, fixed by a cover slip and directly used for microscopy (Olympus BX51). The cell sizes were determined in ImageJ.

## 3. Results

### 3.1. The suppressor mutation in Δ*sll0729* maps to a single nucleotide in the *slr1790* promoter

The absence of M.Ssp6803II resulted in the strongest phenotypic differences among the mutants lacking specific DNA methyltransferase activities, including bluish pigmentation due to decreased chlorophyll levels.^20^ However, the phenotype was not stable, as a relatively high number of suppressor mutants with WT-like pigmentation appeared after prolonged cultivation of mutant Δ*sll0729*.^21^ To identify the mutation(s) leading to reversion of the phenotype, seven independently isolated suppressor clones were sequenced and compared to the parental *Δsll0729* strain. The sequence data revealed several single nucleotide variations (SNV) and short deletions in the different suppressor mutants (**Supplementary Table S1**). However, one SNV was found in the DNA of all suppressor clones. This SNV affected a GGCC motif for M.Ssp6803II-specific DNA methylation changing it to GGTC within the promoter of *slr1790* (**Fig. 1**).

**Figure 1.**
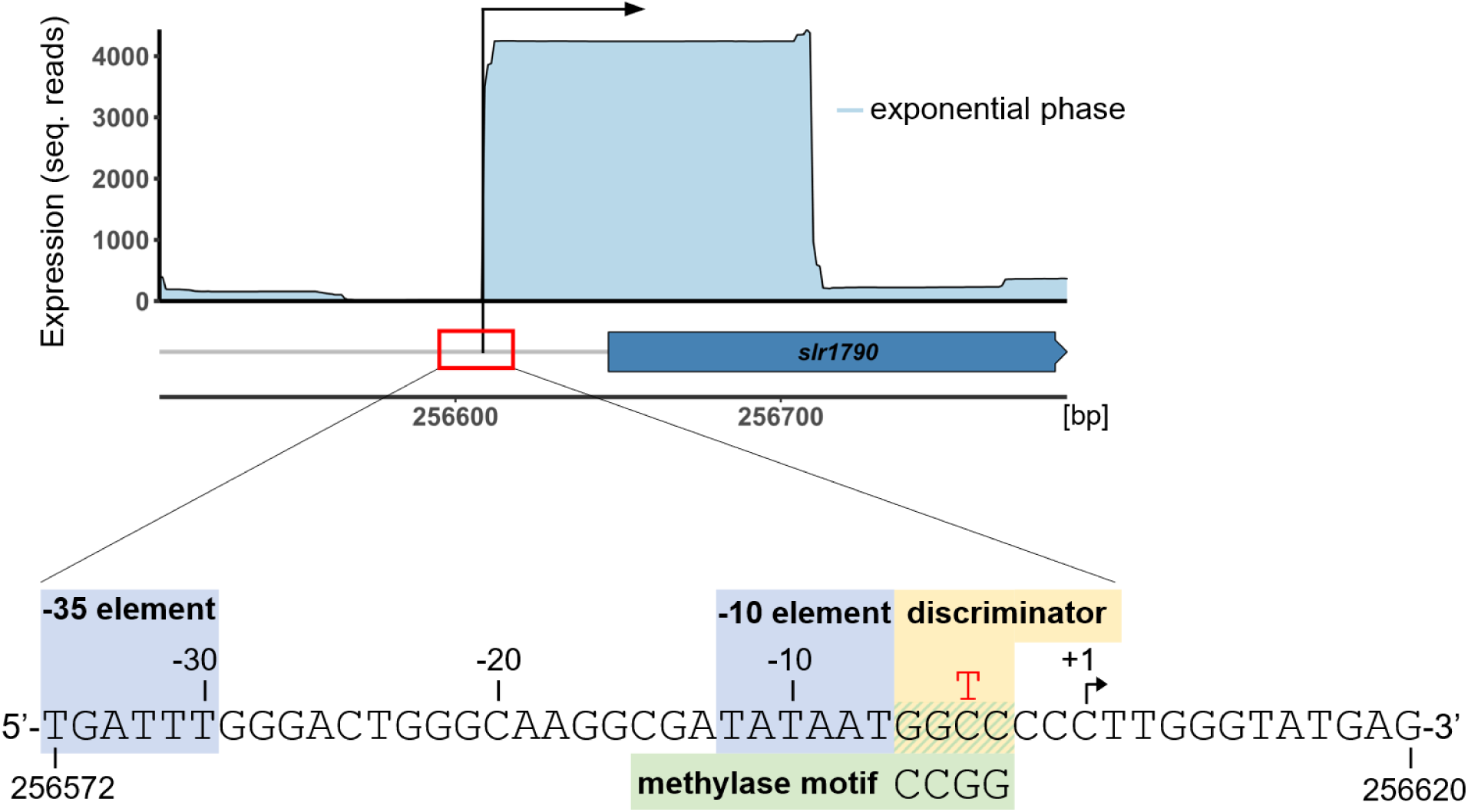
One SNV upstream of *hemJ* is conserved in all suppressor clones. Upper panel: The gene *slr1790* encoding HemJ is transcribed from a single promoter with a transcription start site (arrow, +1) previously mapped to position 256,609 (NCBI reference sequence accession NC_000911.1) of the forward strand and associated with a 5’ UTR of 89 nt. Shown is the accumulation of sequencing reads in the genome-wide mapping of transcription start sites by differential RNA-seq.^31^ The sharp decline in the sequence coverage results from the obtained average sequencing read length of 100 nt.^31^ Lower panel: Resequencing of the genomes of seven suppressor clones revealed one conserved single nucleotide variation (SNV) at position 256,605 of the forward strand (position −4 with regard to the transcription start) changing C to T. This transition changes the GGCC methylation motif to GGTC (red T) and is located between the promoter −10 element and the transcription start site, that is, within the promoter discriminator region (highlighted by the ochre background).

The gene *slr1790* encodes the protoporphyrinogen IX oxidase HemJ.^22,23^ HemJ is essential for the enzymatic protoporphyrin IX (PPIX) synthesis, which is the last common precursor for the biosynthesis of heme and chlorophyll. Heme is not just an essential cofactor of cytochromes and other electron carriers and enzymes, but also a precursor for the synthesis of bilins — the chromophores of phycocyanin and allophycocyanin light harvesting proteins, and of photoreceptors such as cyanobacteriochromes and phytochromes.^32^ Therefore, a possible defect in the expression of a gene necessary for tetrapyrrole biosynthesis could be related to the observed pigmentation phenotype of the Δ*sll0729* mutant.

The conserved SNV in the suppressor mutants at position 256,605 affects specifically the GGCC motif for M.Ssp6803II-specific DNA methylation in the core promoter of *slr1790*. The motif is situated immediately downstream of the −10 element and just upstream of the transcriptional start site (position −4; **Fig. 1**) of this gene. This region is crucial for correct transcription initiation, because DNA strand separation by RNA polymerase in the formation of the open complex is controlled by an arranged separation of specific nucleotides starting from −11 and passing the discriminator sequence.^33^

Overall, there are 38,512 instances of the GGCC motif on one chromosome in the multi-copy *Synechocystis* 6803 genome.^20^ Hence, it is possible that not all of them are fully methylated. Bisulfite sequencing analysis permits the direct and highly sensitive detection of ^m5^C but it can also be used to map ^m4^C, although ^m4^C is partially resistant to bisulfite-mediated deamination. When the assay was used for a global methylation analysis, we found that about 90% of GGCC methylation sites were methylated in the *Synechocystis* 6803 genome. Among them, bisulfite sequencing showed a complete methylation of the GGCC at position 256,605 in the DNA isolated from WT cells (**Supplementary Fig. S1**), which will be absent in mutant Δ*sll0729* with abolished M.Ssp6803II and in suppressor clones bearing the mutated *slr1790* promoter with GGTC.

### 3.2. Lack of GG^m4^CC methylation in *Synechocystis* 6803 impacts *hemJ* expression

We have previously reported that the lacking methylation of GGCC motifs by M.Ssp6803II impacted the expression of genes including *sll0470* and *sll1526*, with log_2_FCs of +1.3 and −1.05, respectively ^21^. Therefore, it appeared possible that the lack of GG^m4^CC methylation had also consequences for the expression of *hemJ*. We reanalyzed our existing transcriptome data sets comparing the gene expression in mutant Δ*sll0729* with WT.^21^ Indeed, a similarly decreased expression of *hemJ* in Δ*sll0729* compared to WT was observed under different light conditions in an experiment from 2013 (**Supplementary Fig. S2A**). In the transcriptome data from 2014, the expression of *hemJ* was diminished in mutant Δ*sll0729* relative to WT (log_2_FC −0.855, below our significance threshold of −1), while in the *sll0729* complementation strain it returned to WT-levels (**Supplementary Fig. S2B**). Finally, in a third dataset from 2017, clearly diminished *hemJ* transcript levels were observed for two freshly generated Δ*sll0729* independent deletion mutants (**Supplementary Fig. S2C**). Thus, these transcriptomic data showed that the expression of *hemJ* was lowered in Δ*sll0729*, in which the methylase M.Ssp6803II specific for cytosine N4-methylation leading to GG^m4^CC was missing. The quantitative difference was in all cases less than twofold compared to WT, but observed reproducibly in three independent experiments performed over several years.

To verify this observation, the expression of *hemJ* was quantified by qRT-PCR in cells of freshly generated Δ*sll0729* mutants and three independently recovered suppressor clones. Again, the absence of M.Ssp6803II resulted in lowered *hemJ* expression in mutant cells, while the expression returned to levels slightly higher than WT in the suppressor clone (**Fig. 2A**). Furthermore, the impact of different point mutations in the discriminator region of the promoter was tested. For this experiment, we fused the reporter genes *luxAB* to the native *hemJ* promoter with intact GGCC motif or variants with a GGAC, GGTC, or GGGC motif (listed in **Supplementary Table S2**). The exchange of GGCC to GGTC (as found in the suppressor clones) resulted in slightly higher transcript levels than the native GGCC motif in the WT background, whereas the GGAC and GGGC variants showed the highest promoter activities (**Fig. 2B**).

**Figure 2.**
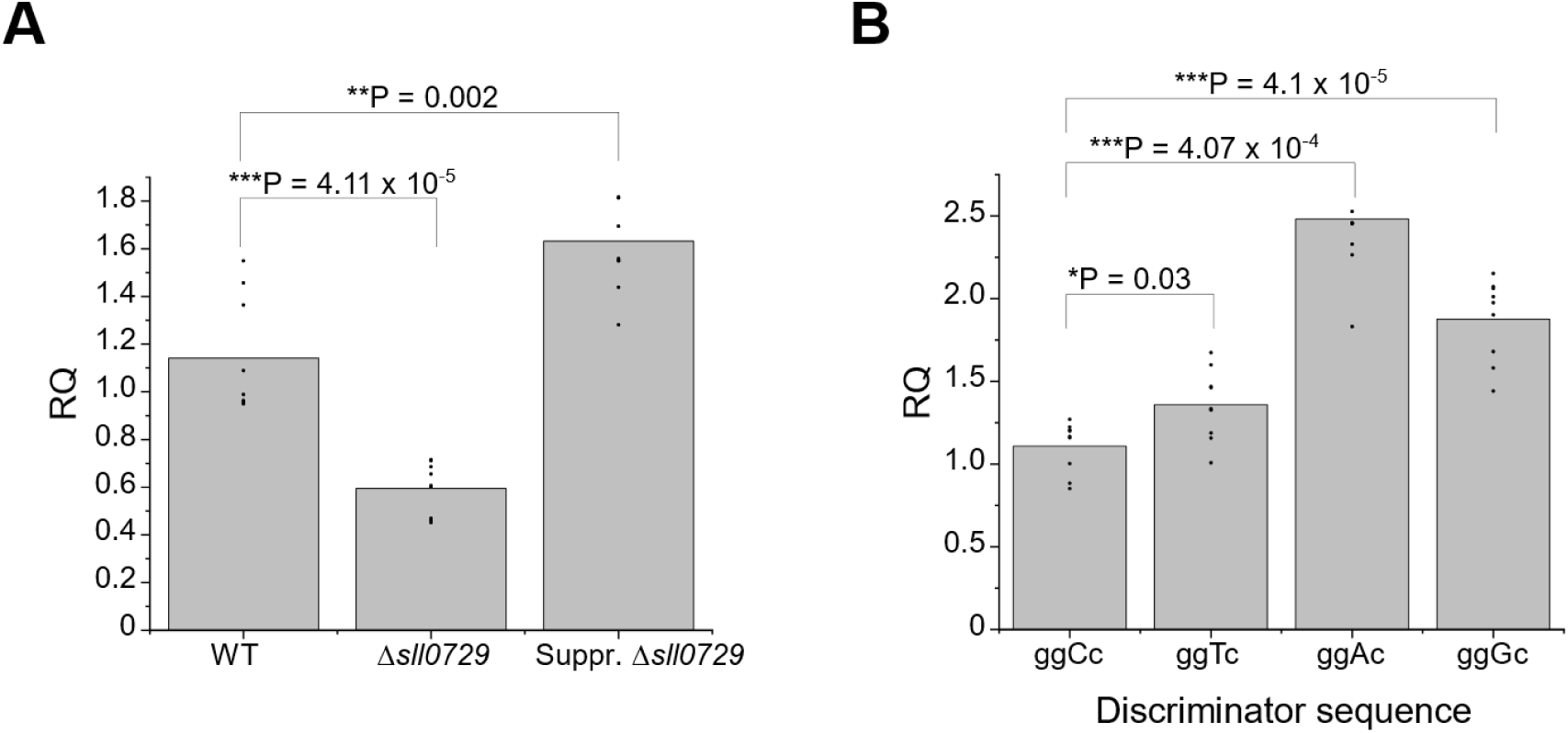
Abundance of transcripts driven by *hemJ* promoter variants in different *Synechocystis* 6803 strains. **(A)** Expression of *hemJ* in wild type (WT), Δ*sll0729*, or one selected suppressor mutant clone (Suppr. Δ*sll0729*) quantified by qRT-PCR. The bars show averages of technical triplicates of biological triplicates, individual data points are given as well. WT was used as the calibrator. (**B)** qRT-PCR quantification of *luxAB* mRNA levels that were under control of *hemJ* promoter variants, in which the first C in the GGCC motif was unchanged or exchanged to A, T or G. The bars indicate averages of technical triplicates of biological triplicates, individual data points are given as dots. The WT motif GGCC was used as the calibrator for Relative Quantification (RQ). Significance was calculated with an unpaired Two-Samples Wilcoxon Test using RStudio; \**P* < 0.05; \*\**P* < 0.01; \*\*\**P* < 0.001) between the strains at corresponding time points (details of analysis and statistical test in **Dataset EV1 and EV2**).

From these experiments we conclude that single nucleotide substitutions in the discriminator region indeed affected the levels of mRNAs produced by the *hemJ* promoter, regardless of whether the native or with the *luxAB* mRNA another transcript was produced. The GGTC motif yielded the quantitatively most similar mRNA level compared to the (methylated) GGCC in the WT background.

### 3.3. Reconstitution of *Synechocystis* 6803 strains to verify the importance of GGCC methylation for *hemJ* expression

The above experiments clearly showed that the identity of the nucleotide at the −4 position in the *hemJ* promoter determined the level of the transcribed mRNA. The measured quantitative differences were relatively minor, but the high frequency of suppressor mutants at this position for the Δ*sll0729* phenotype suggested that these differences were physiologically meaningful. Therefore, we tested whether the GGCC motif and proper expression of *hemJ* were directly related to the phenotypic stability of Δ*sll0729.* Several strains were generated to address this possibility (**Supplementary Table S2**). Attempts to completely segregate the *slr1790* deletion were unsuccessful, consistent with an earlier report.^22^ However, we were able to generate two strains, one with the native *slr1790* promoter (P*slr1790*) and one with the mutated (GGCC substituted by GGTC) *slr1790* promoter (MP*slr1790*) by gene replacement (both strains harbored upstream a spectinomycin resistance gene) (**Supplementary Table S2**). Genotyping and sequence analysis revealed that both strains were completely segregated and stably maintained the respective promoter variants (**Fig. 3A, B**).

**Figure 3.**
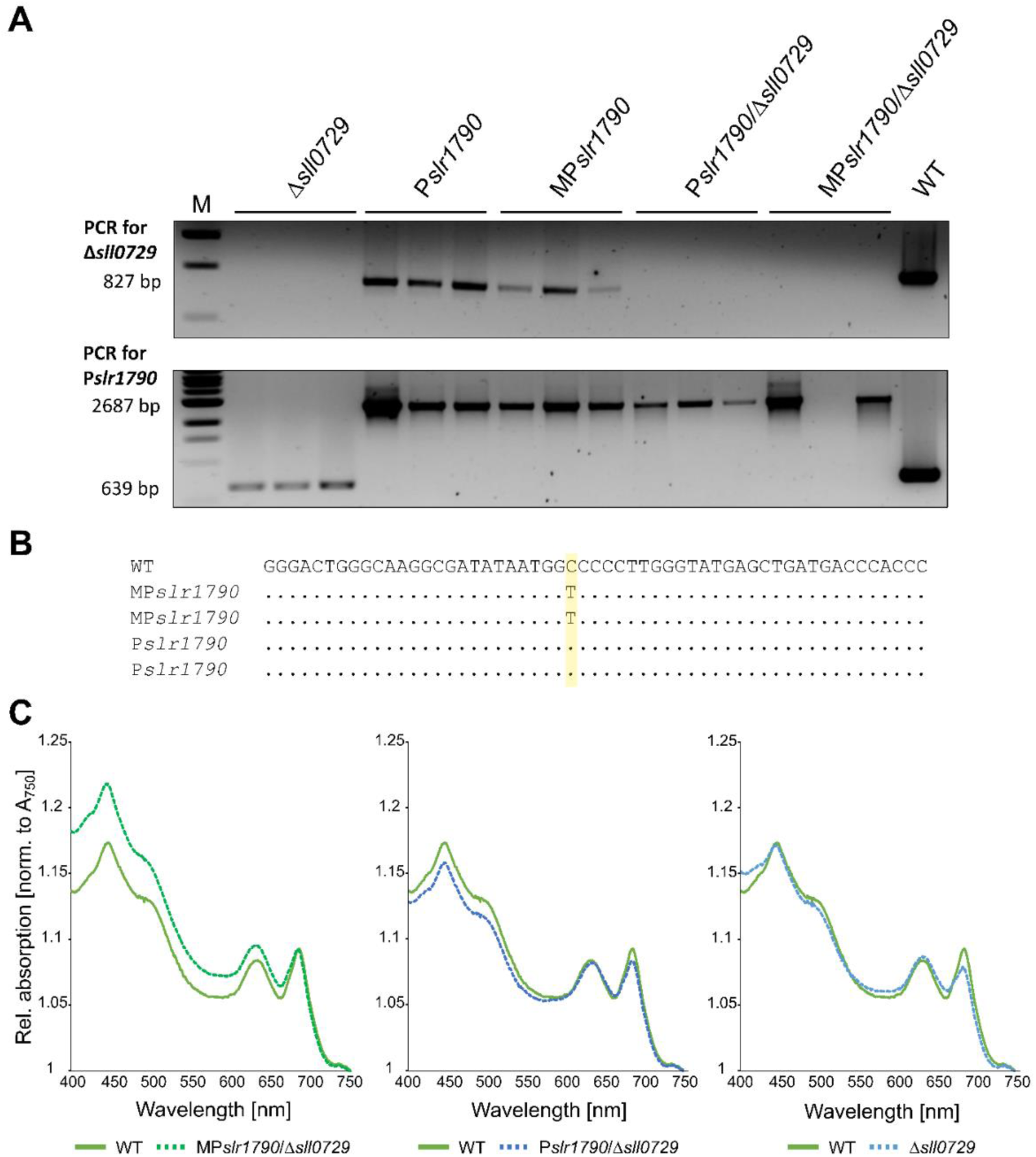
Genotyping and phenotyping of *Synechocystis* 6803 strains to verify the importance of GGCC methylation for *hemJ* expression. (**A**) Genotyping via PCR. Upper panel, the gene encoding M.Ssp6803II is not present in strains with Δ*sll0729* background as indicated by the lack of the 827 bp amplicon. Lower panel, presence of the wild-type (WT) or the mutated *hemJ* promoter in the generated strains as indicated by the 639 bp amplicon in WT and Δ*sll0729*, or the 2687 bp fragment in the manipulated strains. The latter fragment is larger due to the *aadA* antibiotic resistance cassette inserted upstream of the promoter. Three independent clones were analyzed per strain. For details of the respective strains, see **Supplementary Table S2**. (**B)** Sequence analysis to verify the intact GGCC motif in the native promoter and its change to GGTC in the clones with mutated promoter sequence. (**C)** Absorption spectra of different strains grown under standard conditions. The spectra were normalized to the absorption at 750 nm.

Subsequently, these two strains and the corresponding WT served as recipients for a new round of *sll0729* deletions, yielding strains Δ*sll0729,* P*slr1790*/Δ*sll0729,* and MP*slr1790*/Δ*sll0729* (**Supplementary Table S2**). The number of kanamycin-resistant clones obtained differed between strains containing either the native or mutated *hemJ* promoter. In the latter case, more clones appeared and these were almost all completely segregated, whereas transformation of the *sll0729* deletion cassette into the WT or the strain harboring the native promoter variant resulted in lower ratios of clones with completely segregated *sll0729* mutation (**Supplementary Fig. S3**).

This finding is in line with our expectations, because the GGTC promoter variant found in the suppressor clones makes the deletion of M.Ssp6803II less harmful for *Synechocystis* 6803. Interestingly, the original bluish phenotype and decreased growth of mutant Δ*sll0729* were observed in freshly segregated clones from transformations of the WT and the strain with the native *hemJ* promoter, while the strain with the mutated *hemJ* promoter and segregated *sll0729* showed WT-like pigmentation, as observed for newly selected suppressor clones (**Fig. 3C, Supplementary Fig. S4**).

Subsequently, selected clones were tested in liquid and solid media under different growth conditions. Many features reported by Gärtner et al.^21^ characterizing the initial Δ*sll0729* mutation were reproduced, such as the smaller cell size, changed pigmentation, and growth (**Supplementary Fig. S4, Supplementary Table S3**). WT cells displayed an average area of 4.80 µm^2^, whereas all cells of strains with deleted M.Ssp6803II were smaller (Δ*sll0729*: 3.91 µm^2^; P*slr1790*/Δ*sll0729*: 3.52 µm^2^; MP*slr1790*/Δ*sll0729*: 3.02 µm^2^). Control strains had slightly larger cell sizes (P*slr1790*: 6.00 µm^2^; MP*slr1790*: 5.51 µm^2^). One of the most obvious changes was related to pigmentation. As previously reported,^21^ fresh mutants lacking M.Ssp6803II had reduced chlorophyll levels if *hemJ* was driven by its native promoter. In contrast, the GGCC-->GGTC mutation of the *hemJ* promoter prior to deleting *sll0729* yielded WT-like pigmentation, which was also observed in suppressor strains. Moreover, segregated Δ*sll0729* mutants in strains with the native *hemJ* promoter were less frequent, less viable, and showed slower growth than WT or Δ*sll0729* mutants in the strain with the GGTC-changed *hemJ* promoter (**Supplementary Fig. S5**).

Most importantly, all clones with a completely deleted *sll0729* gene were not further able to methylate GGCC sites, regardless of the strain from which the DNA was isolated. In contrast, DNA from all strains with an intact *sll0729* gene was resistant to restriction enzymes that cut only DNA with non-methylated GGCC motifs (**Supplementary Fig. S6**). Finally, in the suspension of segregated fresh Δ*sll0729* and P*slr1790*/Δ*sll0729* mutant clones with bluish phenotype, clones with WT-like pigmentation spontaneously appeared at a high frequency after one to four weeks of cultivation, as observed previously. DNA was isolated from two of these putative suppressor clones and the *hemJ* promoter region was PCR-amplified and sequenced. As observed in the genomic analysis of suppressor lines, the GGCC motif upstream of the *hemJ* transcription start site was again changed at position −4 to GGTC.

### 3.4. Effects of epigenetic manipulation on pigmentation

The gene *slr1790* encodes protoporphyrinogen IX oxidase PPO (HemJ),^22,23^ the enzyme converting the colorless protoporphyrinogen IX (PPIX) into red PPIX (**Fig. 4**). This is the last common step in the biosynthesis of heme and chlorophyll cofactors and of all further tetrapyrrole derivatives ^32^. Therefore, the relatively small differences in *hemJ* transcription caused by the nature of the nucleotide at the −4 position of the promoter might directly affect the formation of pigment biosynthesis intermediates.

**Figure 4.**
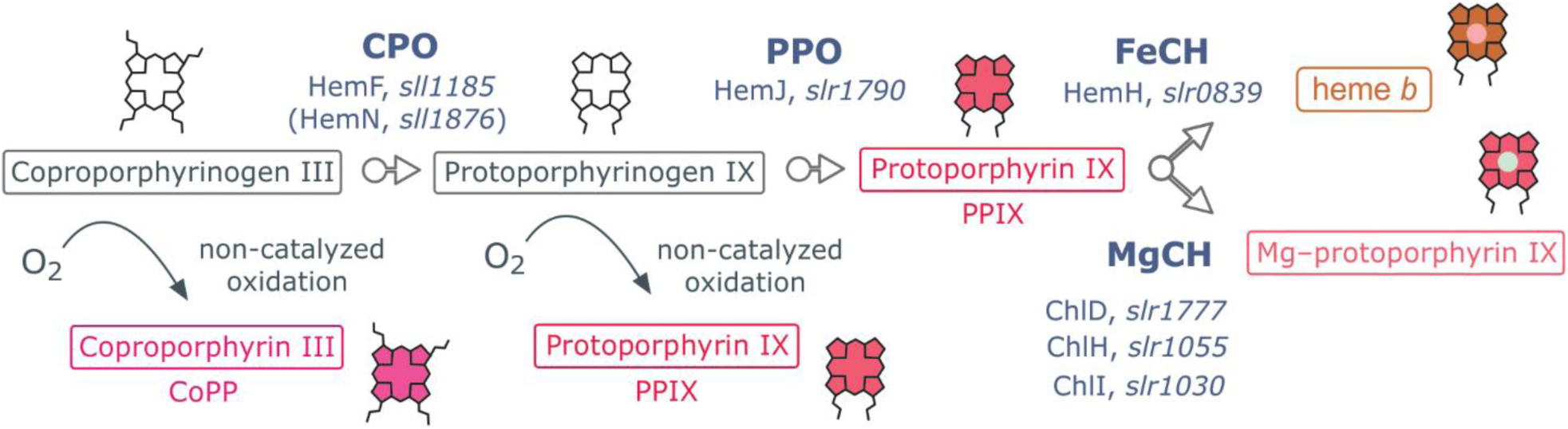
HemJ catalyzes the enzymatic conversion of protoporphyrinogen IX to PPIX. Scheme of last common steps of the chlorophyll and heme biosynthesis. HemJ is one of three different paralogs of the protoporphyrinogen IX oxidase (PPO) known in nature and is present in most cyanobacteria such as *Synechocystis* 6803. This enzyme oxidizes protoporphyrinogen IX into PPIX, the universal precursor of chlorophylls, hemes and bilins ^22,23^. The enzyme names and accession numbers of the corresponding genes in *Synechocystis* 6803 are given.

To obtain a better picture of chlorophyll biosynthesis, intermediates in porphyrin biosynthesis leading to chlorophyll were quantified in different strains grown under identical conditions (**Table 1**). The most remarkable differences were found in the amounts of CoPP and PPIX. CoPP is the non-enzymatically oxidized form of coproporphyrinogen III, which is the substrate of coproporphyrinogen oxidase, a preceding enzymatic step of HemJ. These two enzymes are enzymatically coupled, and the impaired activity of HemJ also affects the decarboxylation of coproporphyrinogen III.^23^ Although PPIX is the product of the HemJ catalytic activity, it can also accumulate in the cell by spontaneous oxidation of protoporphyrinogen IX after blocking HemJ activity (**Fig. 4**,^23^). Unfortunately, the cellular content of protoporphyrinogen IX cannot be measured directly by HPLC with fluorescence detection, as this compound is colorless, and its non-enzymatic oxidation to PPIX is very fast.

**Table 1.** Quantification of chlorophyll synthesis intermediates in different strains of *Synechocystis* 6803 with mutated *sll0729* gene and/or mutated *slr1790* promoter. The first committed intermediate of chlorophyll pathway is Mg-protoporphyrin IX (MgP), which is consequently methylated (MgPME). The MgPME is converted into divinyl protochlorophyllide (DV-Pchlide) and then, in two steps, into monovinyl chlorophyllide (MV-Chlide); chlorophyll is finally made by attachment of phytol to the MV-Chlide. Relative values (peak areas) from six biological replicates analyzed in two runs are shown. Statistically significant differences (p value ≤ 0.05) are printed in bold. Statistical details are in **Supplementary Tables S5** and **S6**.

|  | CoPP | PPIX | MgP | MgPME | DV-Pchlde | MV-Chlide |
| --- | --- | --- | --- | --- | --- | --- |
| WT | 65.1 | 49.2 | 100.3 | 443.2 | 200.2 | 2328.6 |
| ***Δslr0729*** | **269.7** | **3921.2** | 88.2 | 340.7 | 161.1 | 2359.5 |
| ***Pslr1790*** | 51.5 | 32.4 | 53.6 | 254.6 | 194.6 | 2336.5 |
| **MP*slr1790*** | 56.6 | 26.5 | 46.8 | 242.6 | 182.3 | 1809.9 |
| ***Pslr1790/Δslr0729*** | **430.8** | **3058.4** | 82.2 | 556.2 | 161.4 | 3377.3 |
| **MP*slr1790/Δslr0729*** | 34.8 | 15.3 | 43.7 | 147.4 | 103.5 | 2752.1 |

Extremely high amounts of PPIX, together with high amounts of CoPP were specifically observed in strains showing the bluish phenotype (**Supplementary Fig. S4**), that is, strains with segregated deletion of *sll0729* and native *hemJ* promoter, whereas all other strains contained low amounts of PPIX and CoPP, similar to WT (**Table 1, Supplementary Tables S5** and **S6**).

These results indicate that the lowered *hemJ* expression dramatically impacted the tetrapyrrole biosynthetic pathway leading to the accumulation of the phototoxic intermediates CoPP and PPIX. We also monitored the levels of later intermediates of the chlorophyll biosynthetic pathway, but detected no clear differences, as in the case of CoPP or PPIX. The lower chlorophyll level observed in the *sll0729* mutant (**Fig. 3C**) is thus most likely caused by extremely high PPIX content, causing strong oxidative stress, rather than restricted chlorophyll biosynthesis.

## 4. Discussion

The primary aim of our study was to identify the genetic changes in frequently occurring suppressor clones that exhibited wild-type-like pigmentation in the presence of abolished GG^m4^CC methylation. A similar frequent appearance of suppressor mutations has recently been reported for other *Synechocystis* 6803 mutants affected in the biosynthesis of tetrapyrroles,^34,35^, which reflects the essential role of this process in the highly pigmented cyanobacterial cell. Our genome sequencing of suppressor clones for Δ*sll0729* revealed one common SNV in the GGCC motif immediately downstream of the −10 sequence in the *slr1790* promoter towards GGTC. The absence of GGCC methylation resulted in slightly diminished *slr1790* expression encoding HemJ, which is involved in porphyrin synthesis, while the complementation of M.Ssp6803II activity due to ectopic expression of *sll0729* reversed *slr1790* expression to WT levels.

Overall, our data indicate that the improper expression of *slr1790* due to missing ^m4^C-methylation in the *hemJ* promoter resulted in unbalanced porphyrin biosynthesis, leading to massive accumulation of CoPP and PPIX. These compounds have great potential to produce reactive oxygen species (ROS), which likely led to the observed strongly reduced growth and pigmentation phenotype reported here and previously for *Synechocystis* 6803,^21^ as well as in corresponding algal and plant mutants.^36,37^ We present two main findings, the physiological relevance of a particular GG^m4^CC methylation and the dramatic phenotypic effect of seemingly only slightly disturbed expression of HemJ.

Our data unambiguously show the relevance of GG^m4^CC methylation at a critical promoter position for the transcription of *hemJ* and the proper biosynthesis of essential tetrapyrroles in *Synechocystis* 6803. If this setting evolved as the most physiological promoter configuration, it should also be expected for related bacteria. There are not many cyanobacteria with mapped transcription initiations sites, but for the related *Synechocystis* sp. PCC 6714 (*Synechocystis* 6714) such a dataset exists.^38^ The two strains share 2838 protein-coding genes, but also have 845 (*Synechocystis* 6803) and 895 unique genes (*Synechocystis* 6714)^39^ indicating substantial genetic differences. Interestingly, a GGCC motif is situated at a comparable position between the −10 element and the transcription start site of *hemJ* in *Synechocystis* 6714, as in *Synechocystis* 6803 (**Supplementary Fig. S7**). Moreover, *Synechocystis* 6714 possesses the gene D082_01520 (GenBank accession AIE72681), which encodes a likely ortholog of the DNA methyltransferase M.Ssp6803II (59% identical and 73% similar amino acids). Hence, the potential impact of GGCC methylation on *hemJ* expression likely is conserved beyond the here investigated model strain *Synechocystis* 6803.

The change from C to T is often observed in eukaryotes, where deamination of the C5 methylated cytosine results in thymine, which escapes the DNA repair system. Such a mechanism is unlikely in the case here because, first, the cytosine in the GGCC motif is N4-methylated; second, the suppressor mutation occurred in cells in which the methylation of this motif is abolished due to deletion of *sll0729*. Hence, we assume that the exchange of C with T was related to the fact that thymine carries a methyl group, which might mimic the previous GGCC motif methylation and thereby restores the proper expression of *hemJ*. Both the methyl groups of ^m4^C and thymidine protrude into the major groove, thereby making it accessible for DNA-binding factors (**Fig. 5**).^2,40^ It has been shown that an interaction between the methyl group and certain amino acids via CH···π hydrogen bonds^41^ of the sigma factor could possibly lead to an stabilized open promoter complex or facilitate recognition of the promoter sequence by the sigma factor.^42,43^ Additionally, exchanging C with T maintains a pyrimidine base at this position. Our expression analysis in WT strains bearing a GGCC or GGTC motif containing *hemJ* promoter in front of the *luxAB* reporter genes verified similar promoter activities of a methylated WT promoter and the mutated promoter sequence (**Fig. 2B**).

**Figure 5.**
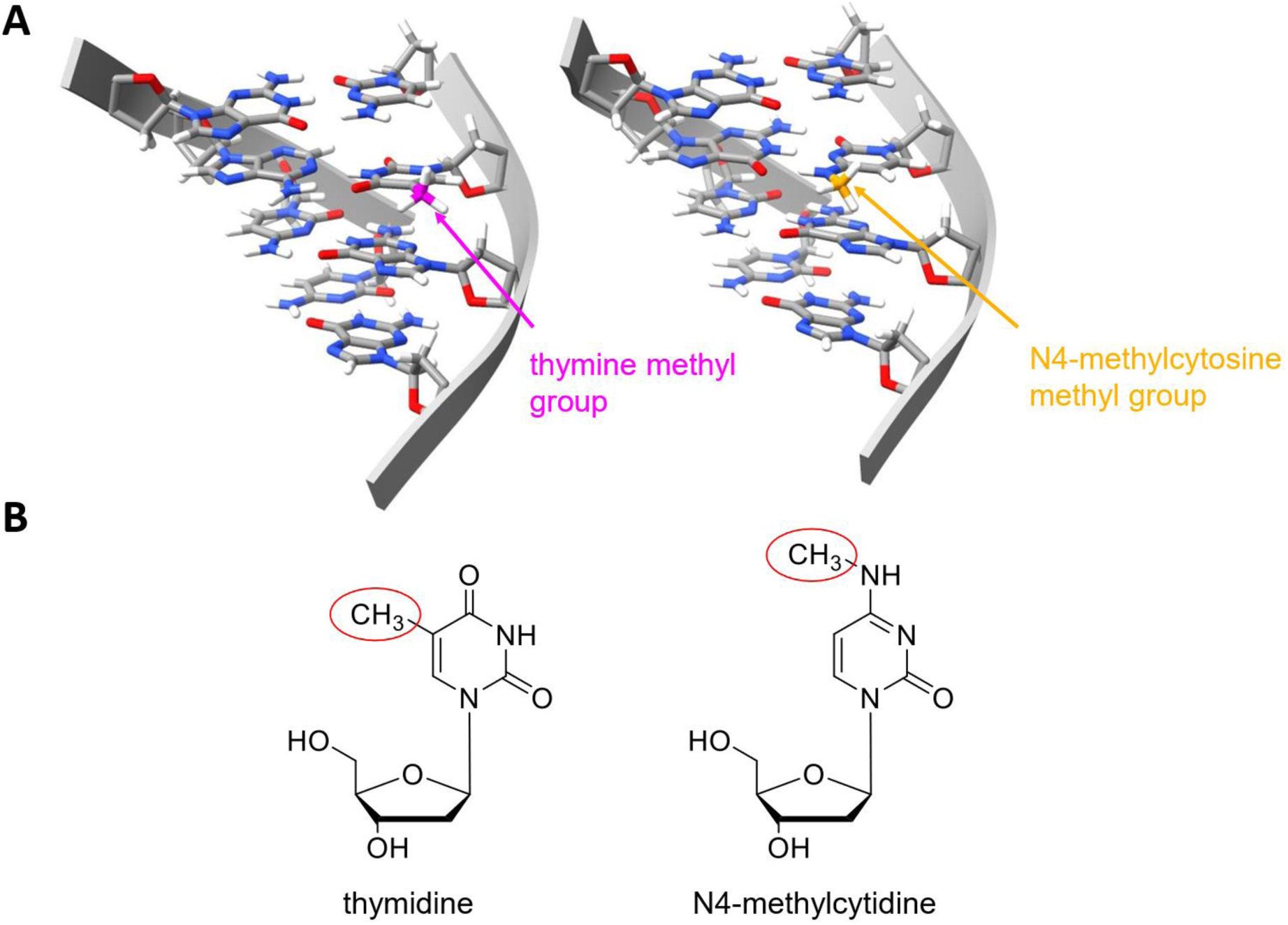
DNA double helix structure of GGC/TC and chemical structures of thymidine and N4-methylcytosine. **(A)** The methyl groups of thymine (purple) and N4-methylcytosine (orange) protrude into the major groove. DNA double helix structure was generated with ChimeraX.^44^ (**B)** Chemical structures of thymidine and N4-methylcytidine. The methyl groups are circled in red. Chemical structures were generated with ChemDraw.

Whether the observed differences in transcription were caused by a direct effect on RNA polymerase kinetics or by a change in transcription factor binding remains unclear. Nevertheless, the GGCC motif methylated by M.Ssp6803II is located in the discriminator sequence of the *hemJ* promoter, which has been shown to have a distinct impact on transcription initiation.^42^ The promoter sequence directs the rate of transcription, starting with RNA polymerase binding, formation of the open promoter complex, and promoter escape, and is crucial for transcription initiation.^45^ Apparent mild changes in this sequence can lead to major changes in transcription initiation.^46–48^ Thus, it is not surprising that the absence of a normally present methyl group at this site can alter gene expression. In another study, hemi-methylation of an adenine located at position −13 in the promoter region of the IS10 transposase gene in the GATC nucleotide sequence led to the activation of transcription.^12^ Methylated bases such as ^m4^C, ^m5^C, and ^m6^A affect the melting properties of the DNA double helix.^49–52^ During open complex formation of the promoter sequence, the melting properties of the DNA double helix lead to altered promoter escape by the RNA polymerase.^33^ In addition, the sequence itself and DNA curvature (bending) should not be neglected, as they could also be affected by the presence or absence of a methyl group.^53^

The second main observation of this study was the strong effect of slightly disturbed *hemJ* expression on the accumulation of certain tetrapyrrole biosynthesis intermediates. This finding is best explained in the context of a multi-enzyme complex for tetrapyrrole biosynthesis that supports the channeling of synthesis intermediates towards the metal chelatases to avoid ROS production, as reported for various organisms.^23,54–56^ The decreased expression of *hemJ* in the absence of GGCC methylation likely led to disturbed complex formation.

Our observation that the level of the first specific chlorophyll precursor, Mg-protoporphyrin IX, was almost unaffected in the Δ*sll0729* strain (**Table 1**) indicates that mutant cells still have sufficient PPIX as a substrate for Mg chelatase. However, protoporphyrinogen IX, released from the complex, probably immediately oxidizes to PPIX (see **Fig. 4**) and initiates ROS production, which then results in further metabolic disturbances and bleaching. Ectopic expression of *hemJ* paralogs in the *hemJ* mutant background of *Synechocystis* 6803 was not successful, likely because of disturbed porphyrin complex formation.^57^ This is another indication that the enzyme-bound conversion of these intermediates in a porphyrin-synthesis complex is necessary for the subsequent correct channeling of PPIX to the insertion of Fe^2+^ or Mg^2+^ by the respective chelatases. A similar ‘bleaching’ phenotype as in *sll0729* deletion strain has been described for a *Synechocystis* ferrochelatase mutant accumulating around 40-times more PPIX than WT. This strain progressively loses chlorophyll despite upregulation of the chlorophyll pathway.^58^

The massive accumulation of the intermediate PPIX in the *sll0729* mutant appears indeed a bit counterintuitive; theoretically, the decreased expression of a key enzyme should result in higher accumulation of its substrate but lower amounts of its products. However, photodynamic herbicides such as acifluorfen, blocking the activity of protoporphyrinogen IX oxidase in plants, have essentially the same effect as the *sll0729* deletion in *Synechocystis* - a dramatic accumulation of PPIX causing severe oxidative damage.^59^ In the Δ*sll0729* strain, this extreme stress situation then initiates high pressure on the cell to solve the problem via improved *hemJ* expression by the mutated GGTC site inside the *hemJ* promoter.

Collectively, our study revealed an important role of ^m4^C-methylation for proper gene expression in *Synechocystis* 6803 and likely other cyanobacteria. Under laboratory conditions, this regulatory feature seems to be particularly important for the regulation of porphyrin biosynthesis, which needs to be tightly regulated to ensure substrate channeling towards metal chelatases to avoid oxidative stress.

^m4^C-DNA methylation is likely to have many more important functions, because highly conserved orthologs of the enzyme M.Ssp6803II are encoded in the genomes of many other cyanobacteria, and many more GGCC motifs exist in the genome sequences of cyanobacteria.^20^ In eukaryotes, epigenetic modifications relevant for gene expression predominantly involve C5-methylcytosine and rarely N6-methyladenine, whereas the here investigated N4-methylcytosine modification is frequent in bacteria. However, recently, the relevance of N4-methylcytosine was discovered as an epigenetic mark in certain invertebrates which use an enzyme acquired from bacteria by horizontal gene transfer.^60^ These findings and the results described here indicate that more attention should be paid to the possible epigenetic effects of ^m4^C-DNA-methylation, both in eukaryotic as well as in prokaryotic organisms.

## Acknowledgments

We thank Klaudia Michl, University of Rostock, for excellent technical assistance and Ingeborg Scholz, University of Freiburg, for the *luxAB* strains. The help of Prof. Klaus Herburger, University of Rostock, during the microscopic analyses of cell sizes is highly appreciated. This study was funded by the German Research Foundation (DFG) (grant HE 2544/12-2 to WRH, grant HA 2002/17-2 to MH) and by project CZ.02.01.01/00/22_008/0004624 of the Czech Ministry of Education, Youth and Sports to RS.

## Data availability

The resequencing data produced in this study are available in the DRA/SRA database with the accession numbers DRR585585–DRR585594, BioProject PRJDB18568. Previously generated bisulfite raw data are available at https://www.ncbi.nlm.nih.gov/biosample/8378604 (BioProject ID: PRJNA430784, BioSample: SAMN08378604, Run: SRX3574087) and previously generated microarray data in the GEO database at https://www.ncbi.nlm.nih.gov/geo/query/acc.cgi?acc=GSE126282, accession number GSE126282 (BioProject PRJNA521475).

## Author contributions

WRH and MH designed the study. NSc and NSt performed most of the experiments and evaluated data. SW and KNM performed resequencing of suppressor clones and data analyses. NSc and RS performed pigment analysis and data analysis. WRH and MH contributed to data analyses. WRH and MH drafted the manuscript. All authors read and approved the final manuscript.

## Supplementary Data

Supplementary data are available at DNARES online.

