## Supplementary Material for "Epigenetic control of tetrapyrrole biosynthesis by ^m4^C DNA methylation in a cyanobacterium"

**Supplementary Datasets:** **p. 2**

**Supplementary Tables:** **p. 3**

**Supplementary Figures:** **p. 9**

**Supplementary References:** **p. 16**

### Supplementary Datasets

#### Supplementary Dataset S1. Details of qRT-PCR analysis and statistical test in Figure 2A.

See separate Excel file.

#### Supplementary Dataset S2. Details of qRT-PCR analysis and statistical test in Figure 2B.

See separate Excel file.

### Supplementary Tables

**Supplementary Table S1. Mutations mapped in the ∆*sll0729* suppressor mutants S1 to S7 (S, strain), compared with the parental ∆*sll0729* strain.** The GenBank (Gbk) accession numbers of the respective replicon is given in the 2nd column, followed by the nucleotide positions (P), the type of mutation (T), with either single nucleotide variation (S) or deletion (D), followed by the respective nucleotides (Nt) and identity of the mutated allele (A), the mutation result (R; dis, promoter discriminator region; fs, frame shift; si, silent), the locus tag (L) of the associated gene, ORF ID (ORF) and gene product.

| S | Gbk | P | T | Nt | A | R | L | ORF | Product |
|---|---|---|---|---|---|---|---|---|---|
| S1, S2, S3, S4, S5, S6, S7 | CP003265 | 256605 | S | C | T | dis | MYO_ 12320 | *slr1790* | protopor-phyrinogen IX oxidase, HemJ |
| S1 | CP003265 | 487987 – 487991 | D | TCCTC | - | fs | MYO_ 14540 | *slr1609* | long-chain-fatty-acid CoA ligase |
| S1 | CP003265 | 703647 | D | A | - | fs | MYO_ 16420 | *slr1393* | cyanobacterio-chrome |
| S3, S4 | CP003265 | 1763737 | S | A | G | Leu > Pro | MYO_ 116270 | *sll1895* | EAL domain protein |
| S2 | CP003265 | 2262055 – 2262059 | D | ATGGT | - | 5’ UTR | MYO_ 120720 | *ssl1807* | membrane protein |
| S5, S6, S7 | CP003265 | 2867061 | S | G | A | si | MYO_ 126030 | *slr0073* | sensory transduction histidine kinase |

**Supplementary Table S2. Overview of genetically modified *Synechocystis* sp. PCC 6803 strains.**

| **Strain** | **Characteristics** | **Resistance** |
|---|---|---|
| P*slr1790* | strain in which native *slr1790* promoter (GGCC motif intact) drives *hemJ* transcription; fused to antibiotic resistance cassette *aadA* upstream; control to MP*slr1790* | spectinomycin |
| MP*slr1790* | mutated *slr1790* promoter (GGCC changed to GGTC) driving *hemJ* transcription and fused to antibiotic resistance cassette *aadA* upstream | spectinomycin |
| Δ*sll0729* | deletion of *sll0729* encoding M.Ssp6803II with inserted antibiotic resistance cassette *aphII* | kanamycin |
| P*slr1790*/Δ*sll0729* | native *slr1790* promoter (GGCC motif intact) driving *hemJ* transcription and fused to antibiotic resistance cassette *aadA* upstream in Δ*sll0729* background | spectinomycin, kanamycin |
| MP*slr1790*/Δ*sll0729* | mutated *slr1790* promoter (GGCC changed to GGTC) driving *hemJ* transcription and fused to antibiotic resistance cassette *aadA* upstream in Δ*sll0729* background | spectinomycin, kanamycin |
| Suppr. Δ*sll0729* | deletion of *sll0729* encoding M.Ssp6803II with inserted antibiotic resistance cassette *aphII* in suppressor strain background | kanamycin |
| P*slr1790* ggCc *luxAB* | Chromosomal integration of *luxAB* with the native promoter *slr1790* in *luxCDE* background | kanamycin, chloramphenicol |
| P*slr1790* ggTc *luxAB* | Chromosomal integration of *luxAB* with GGCC to GGTC mutated *slr1790* promoter in *luxCDE* background | kanamycin, chloramphenicol |
| P*slr1790* ggAc *luxAB* | Chromosomal integration of *luxAB* with GGCC to GGAC mutated *slr1790* promoter in *luxCDE* background | kanamycin, chloramphenicol |
| P*slr1790* ggGc *luxAB* | Chromosomal integration of *luxAB* with GGCC to GGGC mutated *slr1790* promoter in *luxCDE* background | kanamycin, chloramphenicol |

**Supplementary Table S3. Standard deviation and significance of cell size differences.** This table extends the data in **Supplementary Fig. S4.**

| | **Average area [µm$^2$]** | **variance** | **standard deviation** | **p value** |
|---|---|---|---|---|
| **WT** | 4.80 | 1.30 | 1.14 | - |
| **Δ*sll0729*** | 3.91 | 1.11 | 1.05 | 6.18739E-09 |
| **P*slr1790*** | 6.00 | 2.22 | 1.49 | 1.0087E-10 |
| **MP*slr1790*** | 5.51 | 1.67 | 1.29 | 1.03697E-05 |
| **P*slr1790*/Δ*sll0729*** | 3.52 | 0.83 | 0.91 | 5.4733E-17 |
| **MP*slr1790*/Δ*sll0729*** | 3.02 | 0.42 | 0.65 | 6.60528E-31 |

**Supplementary Table S4. Deoxynucleotide primers used in the study.** All sequences are given in 5' to 3' direction; reverse transcriptase (RT).

| Name | Sequence (5' to 3') | Description | Purpose |
|---|---|---|---|
| RT-qPCR_slr1790-rv | ATAGACAAAAAGACGCAC | Primer for Reverse Transcriptase reaction | Quantitative Reverse Transcription PCR |
| RT-qPCR_rnpa-rv | TTAGACGATTTCTAACCGT | Primer for Reverse Transcriptase reaction | |
| RT-qPCR_luxAB-fw | TCTTCCTAACAGGTTAGC | Primer for Reverse Transcriptase reaction | |
| qPCR_slr1790-fw | AAGCGCGAGTATTTCTCC | Primer for qPCR amplification of *slr1790* | |
| qPCR_slr1790-rv | ATCCAGCAAACCAAACAACA | Primer for qPCR amplification of *slr1790* | |
| qPCR_rnpa-fw | CCAGACCGTTTATCAGCAAG | Primer for qPCR amplification of *rnpa* | |
| qPCR_rnpa-rv | TTTTGGCTGACGGTGATG | Primer for qPCR amplification of *rnpa* | |
| qPCR_luxAB-fw | GCAGCAACAAATAAATTTCCCG | Primer for qPCR amplification of *luxAB* | |
| qPCR_luxAB-rv | ATCGCTTTGTTCGGCTTG | Primer for qPCR amplification of *luxAB* | |
| slr1790-AQ-GGCC-fw | AAAAACAAATATTTCCAAACTTCATTTCCCAGGAACAGGGTGGGTC | Aqua Cloning of Promoter *slr1790* WT variant into pILA vector | Generation of luciferase Strains carrying Promoter variants of *slr1790* upstream of *luxAB* |
| slr1790-AQ-GGTC-fw | AAAAACAAATATTTCCAAACTTCATTTCCCAGGAACAGGGTGGGTCATCAGCTCATACCCAAGGGGACCATTATATCGCCTTGC | Aqua Cloning of Promoter *slr1790* GGTC variant into pILA vector | |
| slr1790-AQ-GGGC-fw | AAAAACAAATATTTCCAAACTTCATTTCCCAGGAACAGGGTGGGTCATCAGCTCATACCCAAGGGGCCCATTATATCGCCTTGC | Aqua Cloning of Promoter *slr1790* GGGC variant into pILA vector | |

| | | | |
|---|---|---|---|
| slr1790-AQ-GGAC-fw | AAAAACAAATATTTCCAAACTTCATTTCCCAGGAACAGGGTGGGTCATCAGCTCATACCCAAGGGGTCCATTATATCGCCTTGC | Aqua Cloning of Promoter *slr1790* GGAC variant into pILA vector | |
| slr1790-AQ-rv | GGGATCCAATTGGCAGTGCAGGTCGATACACTCCCTTGGTTTTCACCATTGCC | Aqua Cloning of Promoter *slr1790* | |
| slr1790 Mufw_190 | ACGGTTTCTTCCGCTATGAC | Cloning of *slr1790* | Generation of promoter variants of *slr1790* with 3' spectinomycin resistance |
| slr1790 Murv_1297 | CAACAGTAACGCCACAAAGG | Cloning of *slr1790*/segregation of Promoter *slr1790_aadA* | |
| Promoter_Mut-P | GGACTGGGCAAGGCGATATAATGGTCCCCTTGGGTATGAGCTGATGACC | Mutagenesis of Promoter *slr1790* GGCC to GGTC | |
| Sm fw | ACGAACCCAGTGGACATAAG | Spectinomycin resistence cassette | |
| Sm rev | TCAGGAACCGGATCAAAGAG | Spectinomycin resistence cassette | |
| slr1790 Murv_2502 | AGCTTTCGAGTGCCATTGAC | Sequencing of Promoter *slr1790* GGCC/GGTC motif | |
| Pslr1790_121rev | GGACTTGGTTTAACCCTCCAAATC | Segregation of *slr1790_aadA* | |
| del0729:0729fw | GGGGAAATAAATAAATCAGC | Segregation of *sll0729_aphII* | Generation of *sll0729* knockout strains |
| del0729:0729rev | AACTCTTTACCTTTGGAAGC | Segregation of *sll0729_aphII* | |

**Supplementary Table S5. Standard deviation and significance of PPIX values.** These data extend the results shown in **Table 1**.

| | **PPIX** | **variance** | **standard dev.** | **p-value** |
|---|---|---|---|---|
| **WT** | 49.2 | 2111.2 | 45.9 | - |
| **Δ*sll0729*** | 3921.2 | 4635178.1 | 2152.9 | 0.017 |
| **P*slr1790*** | 32.4 | 636.3 | 25.2 | 0.355 |
| **MP*slr1790*** | 26.5 | 499.7 | 22.4 | 0.222 |
| **P*slr1790*/Δ*sll0729*** | 3058.4 | 1865476.3 | 1365.8 | 3.52E-07 |
| **MP*slr1790*/Δ*sll0729*** | 15.3 | 316.5 | 17.8 | 0.078 |

**Supplementary Table S6. Standard deviation and significance of CoPP values.** These data extend the results shown in **Table 1**.

| | **CoPP** | **variance** | **standard dev.** | **p-value** |
|---|---|---|---|---|
| **WT** | 65.1 | 388.14 | 19.70 | - |
| **Δ*sll0729*** | 269.7 | 71077.08 | 266.60 | 0.007 |
| **P*slr1790*** | 51.5 | 637.81 | 25.26 | 0.085 |
| **MP*slr1790*** | 56.6 | 1423.01 | 37.72 | 0.266 |
| **P*slr1790*/Δ*sll0729*** | 430.8 | 74011.41 | 272.05 | 5.58061E-05 |
| **MP*slr1790*/Δ*sll0729*** | 34.8 | 434.50 | 37.723 | 0.003 |

### Supplementary Figures

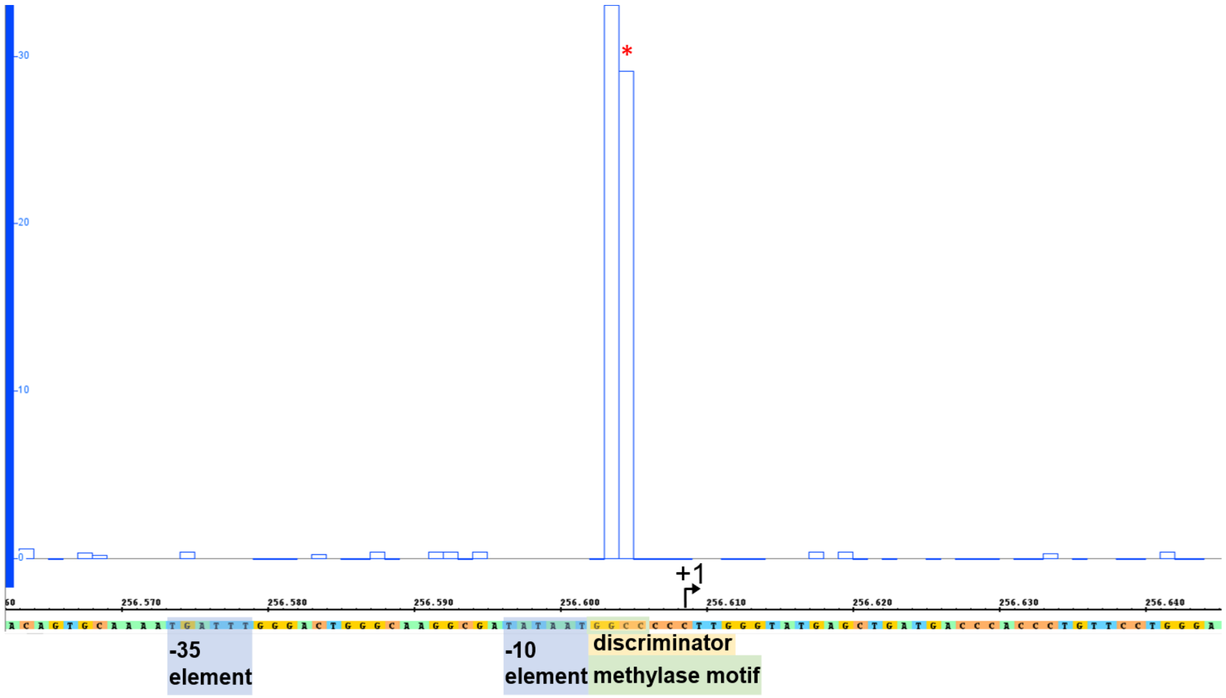


**Supplementary Figure S1. The GGCC-motif in the *slr1790* promoter is fully methylated in wild-type DNA.** Bisulfite sequencing data revealed that the C at pos. 256,605 (marked by red asterisk) is methylated in wild-type cells. Transcriptional start site is indicated as a black arrow (+1). Previously generated bisulfite raw data are available at https://www.ncbi.nlm.nih.gov/biosample/8378604 (BioProject ID: PRJNA430784, BioSample: SAMN08378604, Run: SRX3574087).

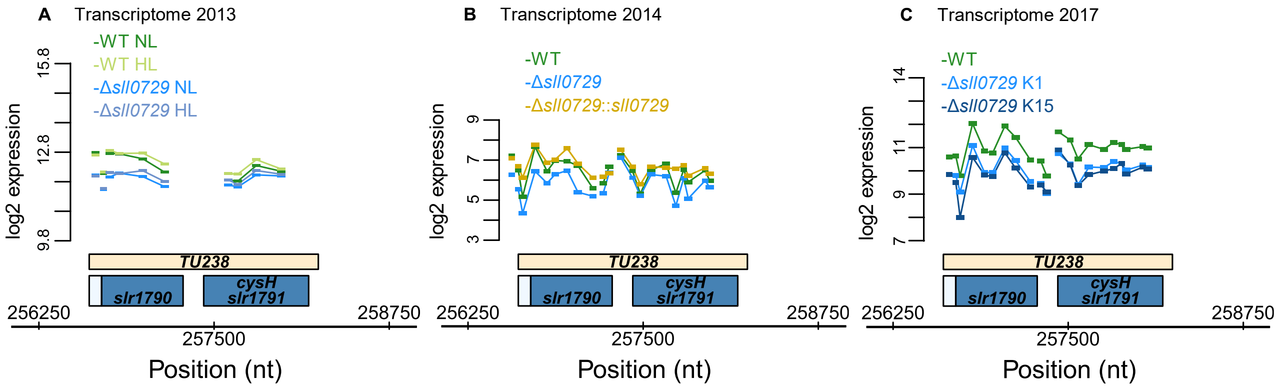


**Supplementary Figure S2. The absence of M.Ssp6803II and connected GGCC methylation/unmethylation impacts *hemJ* expression.** Visualization of the microarray results obtained for the *hemJ-cysH* (*slr1790-slr1791*) locus from three previous analyses with different cultivation schemes and microarray designs. In all analyses, samples from wild type (WT) were compared to an Δ*sll0729* mutant. The two genes are transcribed in a joint transcriptional unit, TU238.[1] **(A)** Experiment performed in 2013 that included a shift from standard light conditions (NL) to high light (HL). (**B)** Experiment performed in 2014 with a freshly made Δ*sll0729* mutant and a strain in which an intact *sll0729* allele was expressed for complementation (strain Δ*sll0729::sll0729*). Note that the array design differed in from that one used in panel (A). Results from this analysis were also used in the publication Gärtner et al. (2019).[2] (**C)** Experiment performed in 2017, same array design as in panel (B), but with two independently obtained new Δ*sll0729* deletion mutants (Δ*sll0729* K1 and K15).

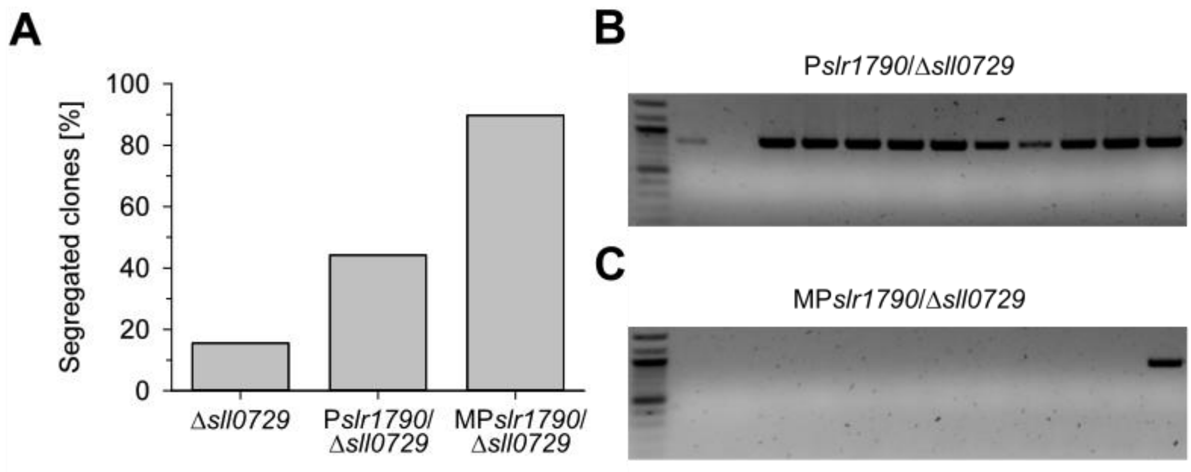


**Supplementary Figure S3. Ratio of successful segregation of the *sll0729* mutation in different *Synechocystis* 6803 strains.** The construct for the deletion of *sll0729* was transformed into cells of the *Synechocystis* 6803 wild type (WT) leading to mutant Δ*sll0729*, the strain with the native (P*slr1790*), and with the mutated *hemJ* promoter (MP*slr1790*). Kanamycin-resistant clones were isolated and the segregation status of the *sll0729* mutation was analyzed via PCR. (**A)** The percentage of completely segregated clones, i.e. where the gene *sll0729* was completely deleted, is displayed. (**B, C)** PCR shows a 823 bp fragment for non-segregated clones of P*slr1790*/Δ*sll0729* and MP*slr1790*/Δ*sll0729*. Only few clones of P*slr1790*/Δ*sll0729* were segregated; however, nearly all MP*slr1790*/Δ*sll0729* were segregated.

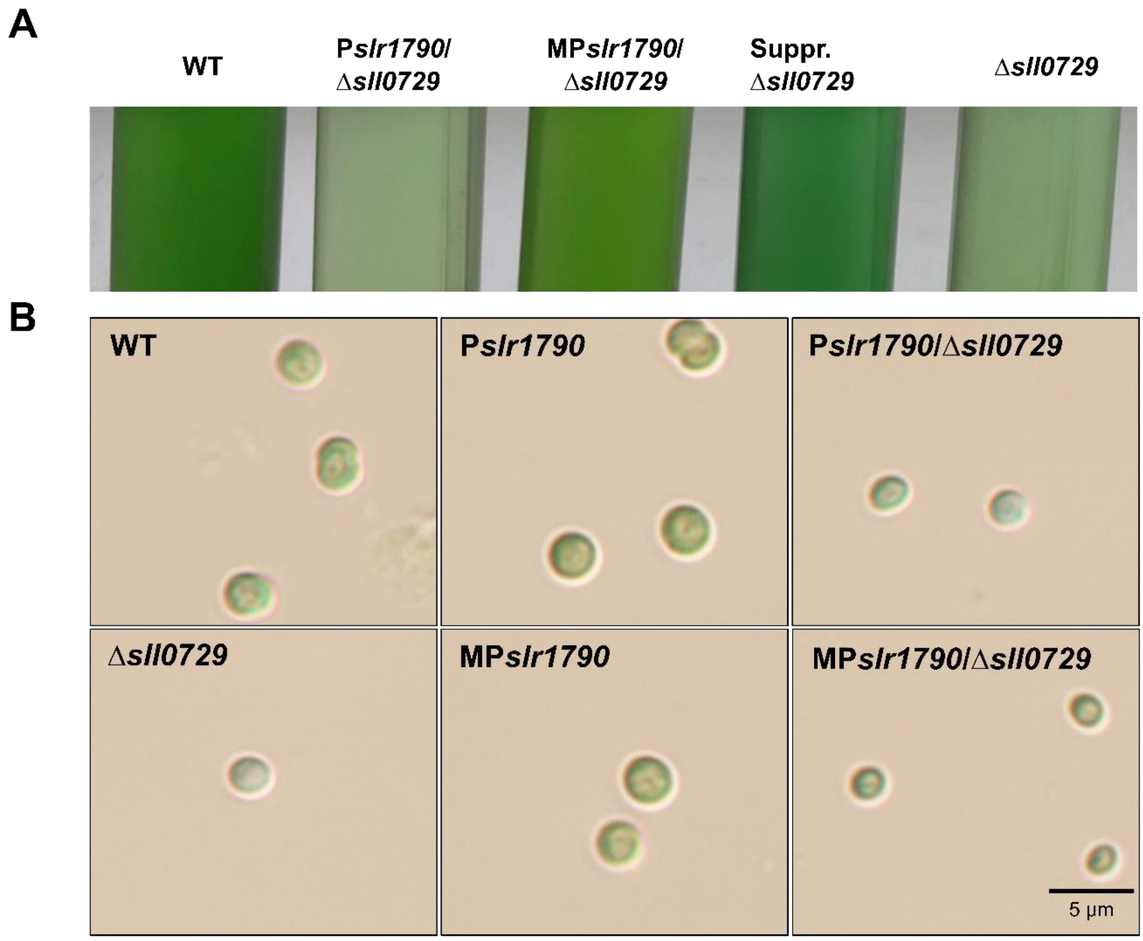


**Supplementary Figure S4. Phenotypic alterations in different strains of *Synechocystis* 6803 with mutated *sll0729* gene and/or mutated *hemJ* promoter.** (**A**) Microscopy images from strain with deleted Δ*sll0729* and *hemJ* promoter mutants. The images where taken at 100x magnification. Please note the different pigmentation of single cells in panel (**B**) Phenotypes, i.e. optical appearance of the different strains. Note the bluish, less pigmented appearance of the mutant Δ*sll0729* and the Δ*sll0729* in the background of the native promoter (P*slr1790*), whereas the pigmentation became WT-like in the suppressor clone (Suppr. Δ*sll0729*) and the strain with mutated *sll0729* gene in the background of the mutated *hemJ* promoter (MP*slr1790*). See **Supplementary Table S3** for numerical values and statistical analysis.

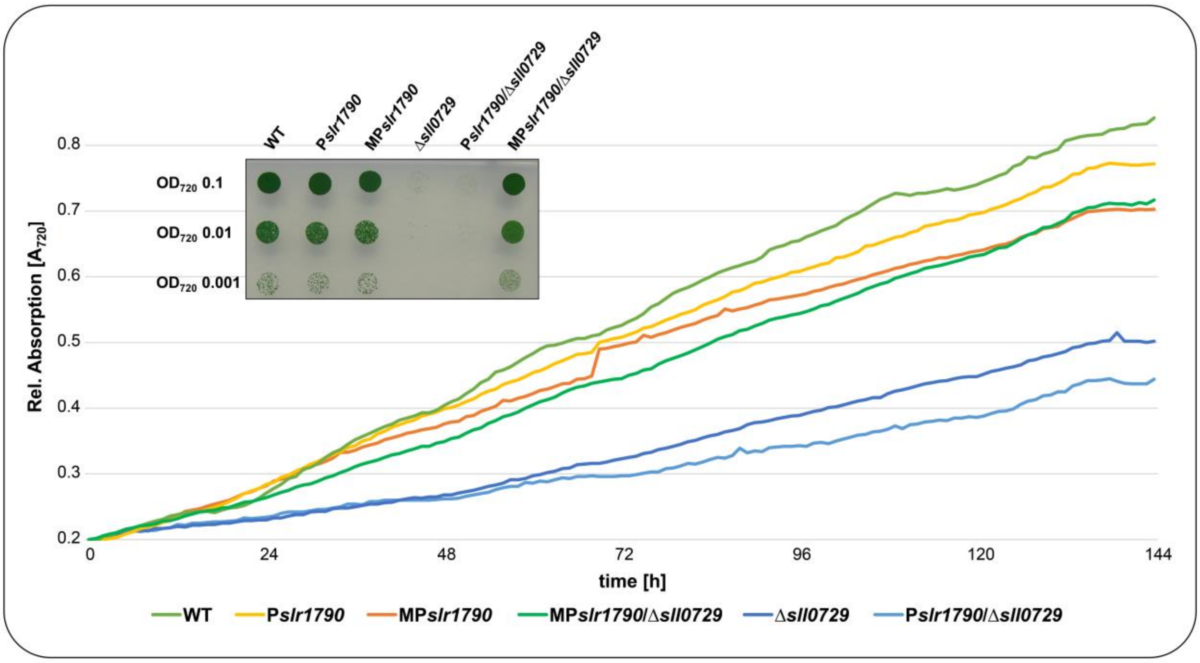


**Supplementary Figure S5.** Growth of different strains of *Synechocystis* 6803 with mutated *sll0729* gene and/or mutated *hemJ* promoter on solid (inset) or in liquid medium. The strain details can be found in **Supplementary Table S2**.

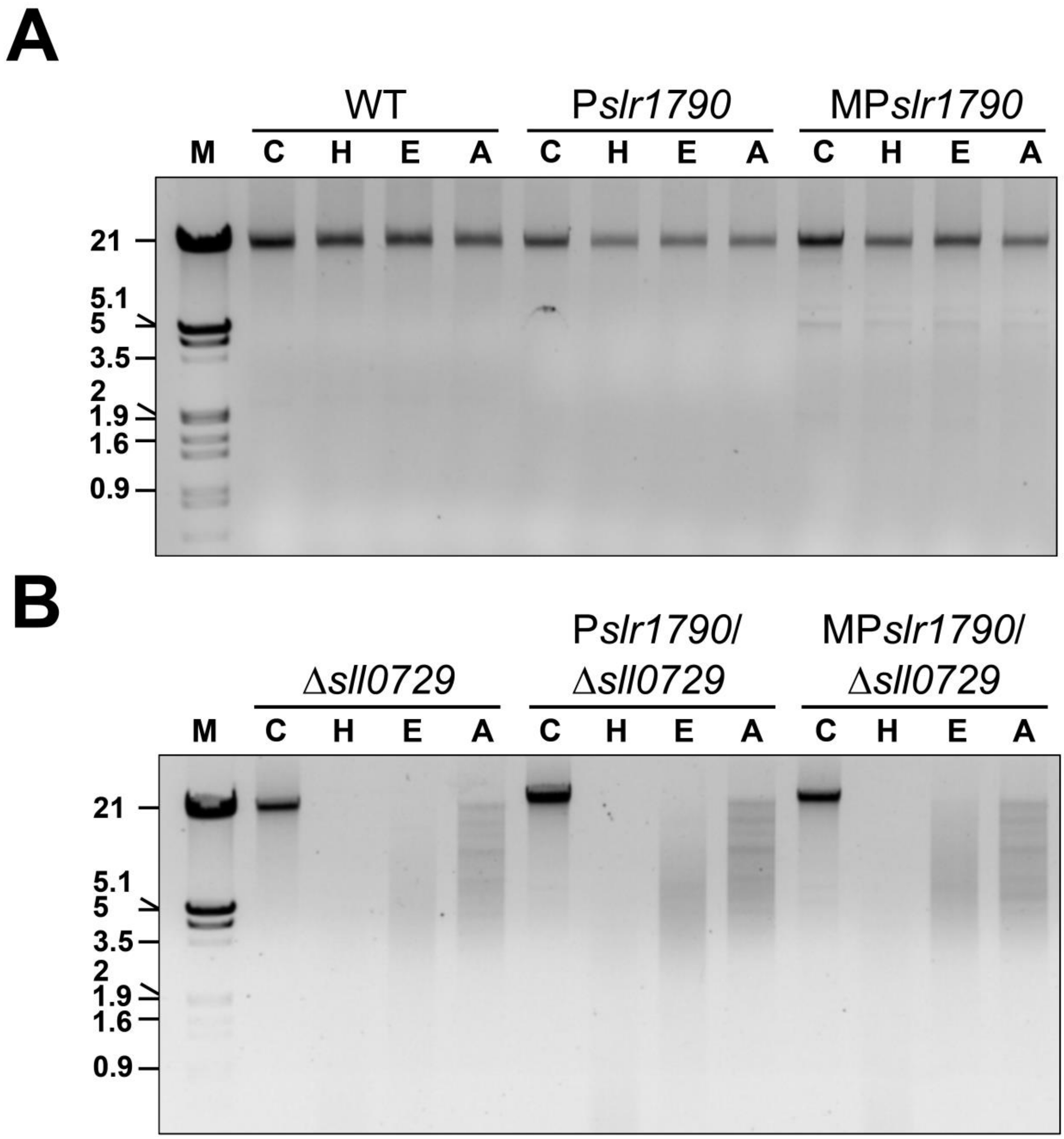


**Supplementary Figure S6. Assay for GGCC methylation of different *Synechocystis* 6803 strains by restriction analysis. (A)** Strains with intact *sll0729* gene expressing M.Ssp6803II. (**B)** Strains with deleted *sll0729* gene not expressing M.Ssp6803II. Three µg total DNA from each strain was treated by *Hae*III (GG/CC), *Eae*I (Y/GGCCR), or *Apa*I (GGGCC/C) for 16 h at 37°C.

These three restriction enzymes bear GGCC motifs in their recognition sequences but cannot cut DNA with methylated GG$^{m4}$CC sites. (M = Marker, combination of bacteriophage λ DNA cut with *Eco*RI and separately with *Hind*III; C = uncut control; H = *Hae*III; E = *Eae*I; A = *Apa*I).

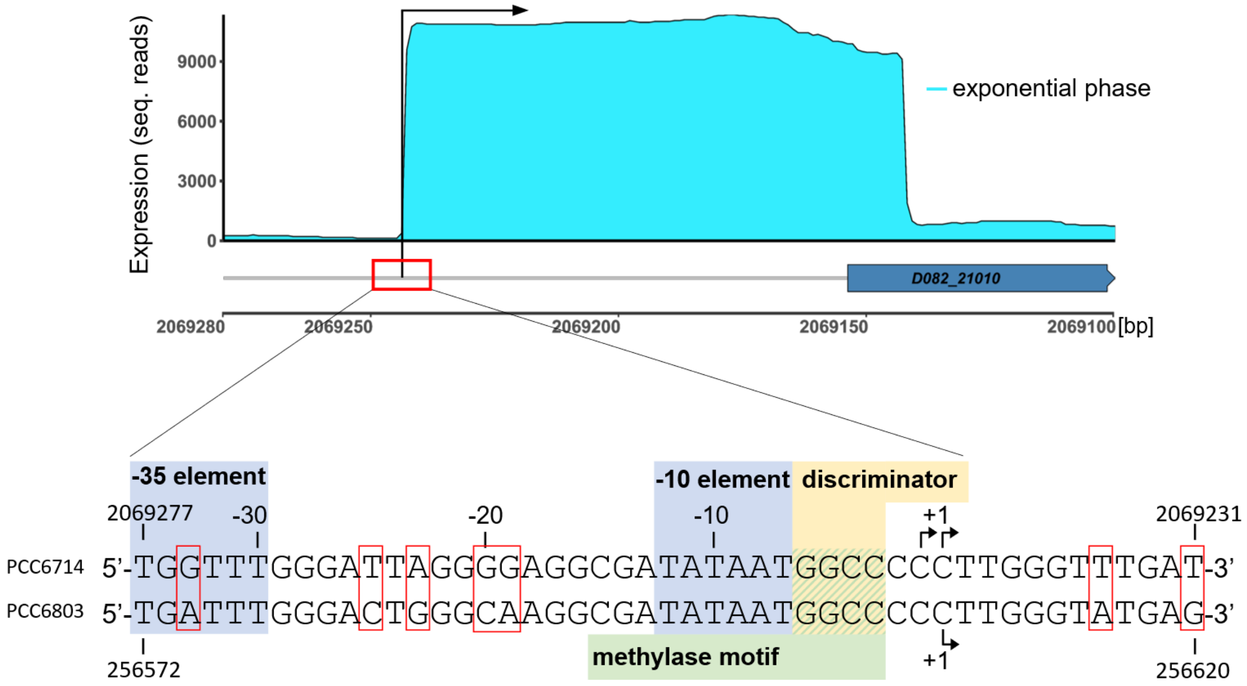


**Supplementary Figure S7. The regulation of *hemJ* via GGCC-specific methylation might be conserved among cyanobacteria.** In the related strain *Synechocystis* 6714 a GGCC-methylation motif is situated between the -10 promoter element and the transcription start site of gene D082-21010 that encodes the HemJ protein (upper sequence).

For comparison, the *Synechocystis* 6803 *slr1790* promoter sequence is included (lower sequence). Sequence differences are boxed in red.
